# Dynamic electrocortical states and paradoxical complexity during desflurane anesthesia

**DOI:** 10.1101/2025.10.13.682019

**Authors:** Duan Li, Anthony G. Hudetz

## Abstract

**Background:** How general anesthesia alters the dynamics of electrocortical activity is crucial to understand the neural mechanisms of unconsciousness. Local cortical activity undergoes spontaneous transitions at constant anesthetic concentration. The spatial organization and temporal dynamics of state transitions in large-scale electrocortical activity is incompletely understood.

**Methods:** Epidural electrocorticogram was recorded from the right hemisphere in 8 rats (14 experiments) using chronically implanted 32-channel flexible electrode arrays during desflurane anesthesia at 6%, 4%, 2%, 0% inhaled concentrations, each maintained for 1 hour. Cortical states were identified by principal component analysis of power spectrograms followed by density-based clustering simultaneously across all anesthetic conditions. State-specific spatiotemporal complexity was quantified by the normalized Lempel-Ziv algorithm to capture signal variability beyond spectral effects. Temporal dynamics were assessed by state occurrence, dwell times, and transition probabilities.

**Results:** Seven cortical states were identified. Six states generally tracked anesthetic depth with an increase in delta power and decrease in complexity, but their occurrence was not tied to any anesthetic level (*p*<0.001). The 7^th^ state was a paradoxical, activated state that mostly occurred during deep anesthesia and was marked by reduced delta (*p*<0.001) and elevated complexity (*p*<0.001). Cortical activity was more likely to remain in a given state than to switch (mean dwell time of 136.55 s, persistence likelihood of 99.36%). When transitions occurred, they followed structured, non-random dynamics, primarily within light- or deep-anesthesia states (FDR-corrected *p*<0.05), with a mild tendency to exit deep states and enter light states (*p*=0.0039), consistent with anesthetic emergence.

**Conclusions:** Electrocortical activity is not a unitary function of anesthetic concentration but involves spontaneous dynamics and paradoxically activated states with increased global complexity during deep anesthesia. The results suggest ongoing spontaneous reorganization of cortical activity during prolonged anesthetic challenge and provide new insight into anesthesia-induced brain dynamics that may inform future strategies for monitoring and manipulating the state of consciousness and facilitating recovery from general anesthesia.

## Introduction

General anesthetics are widely used in clinical practice, yet the neuronal mechanism by which they suppress consciousness is incompletely understood. Traditionally, general anesthesia has been viewed as a unitary function in which brain activity is deterministically tied to anesthetic concentration. However, recent evidence indicates that cortical activity can spontaneously fluctuate between discrete states even at a constant anesthetic level. These states may occur across multiple concentrations, suggesting a complex, many-to-many relationship between cortical dynamics and anesthetic concentrations (Hudson, Calderon et al. 2014, Lee, Wang et al. 2020, Li and Hudetz 2025). Understanding these relationships is crucial for elucidating the neural mechanisms of unconsciousness and for guiding safe, effective clinical anesthesia (Hudson 2017).

Most previous studies reported spontaneous state transitions in localized cortical activity under steady anesthetic levels (Hudson, Calderon et al. 2014, Lee, Wang et al. 2020, Li and Hudetz 2025). For example, in the rat visual cortex, neuronal activity was observed to transition among five states including one that appeared paradoxical: despite occurring under deep anesthesia, it exhibited high, asynchronous spike activity and increased electromyographic signals, suggestive of partial arousal (Lee, Wang et al. 2020). Subsequent analysis, however, revealed relatively low neuronal complexity, suggesting that the paradoxical state was likely an unconscious state (Li and Hudetz 2025). Nevertheless, the generality of these findings remains uncertain, as unit recordings were limited to the visual cortex and thus did not capture large-scale cortical dynamics presumed to be more closely linked to altered states of consciousness.

Previous studies that examined spatial organization and temporal coordination of cortical state transitions under anesthesia have been limited. They typically examined large-scale activity either over short durations at multiple anesthetic concentrations (Curic, Ashby et al. 2024) or under a single steady level (Li, Vlisides et al. 2019) without maintaining prolonged, pharmacologically stable periods. Moreover, although cortical complexity has been proposed as an promising indicator of the state of consciousness (Sarasso, Casali et al. 2021, Li, Fabus et al. 2022), it has rarely been quantified across multiple cortical states (Li and Mashour 2019), leaving questions open about the organization and functional capacity of the anesthetized brain.

To address these gaps of knowledge, we analyzed hemispheric electrocorticography signals in rats using an anesthetic protocol spanning desflurane concentrations of 6%, 4%, 2%, 0%, each maintained for 1 hour. We applied unsupervised clustering to identify brain states independent of anesthetic concentration. We hypothesized that cortical states derived from the spectral properties of electrocorticographic signals would exhibit many-to-many relationships with anesthetic concentrations and would show distinct complexity features associated with state transitions. We further expected to identify a paradoxical state under deep anesthesia, but with low complexity consistent with an unconscious condition. Contrary to our hypothesis, this paradoxical state exhibited high complexity, revealing an unexpected cortical reorganization and a theoretical possibility for some form of conscious processing during prolonged deep anesthesia.

## Materials and Methods

### Experimental Design

The study was approved by the Institutional Animal Care and Use Committee (IACUC) of the University of Michigan. All procedures were performed in accordance with the Guide for the Care and Use of Laboratory Animals of the Governing Board of the National Research Council.

Electrocorticography signals were recorded from the right hemisphere of rats using chronically implanted 32-channel flexible polymer electrode arrays arranged in a 4×8 grid (Neuromicrosystems, Budapest, Hungary) (**Figure 1A**). Part of the cranium was lifted, the electrode was laid over the dura, and the cranial flap was closed and sealed with dental cement as previously described (Fedor, Zatonyi et al. 2020). Eight rats were included in the study, and each underwent two to three experiments for economic reasons and to assess test-retest reliability. In a subset of four rats, an electromyogram electrode was inserted into a nuchal muscle to monitor autonomic arousal as an indicator of impending recovery of consciousness. One to eight days after surgery, animals were placed in a ventilated chamber allowing free movement. Desflurane was administrated at stepwise decreasing concentrations of 6 %, 4 %, 2 %, and 0 % vaporized into 30% O_2_, balance N_2_ (**Figure 1B**). Anesthetic concentration was continuously monitored (POET IQ2, Criticare Systems, Inc., Waukesha, WI, USA). Core body temperature was maintained at 37 °C using subfloor heating. A 15-min equilibration period was allowed between successive anesthetic levels before the electrophysiological recordings. Spontaneous electrocorticographic activity was recorded for one hour at each concentration.

**Figure 1.**
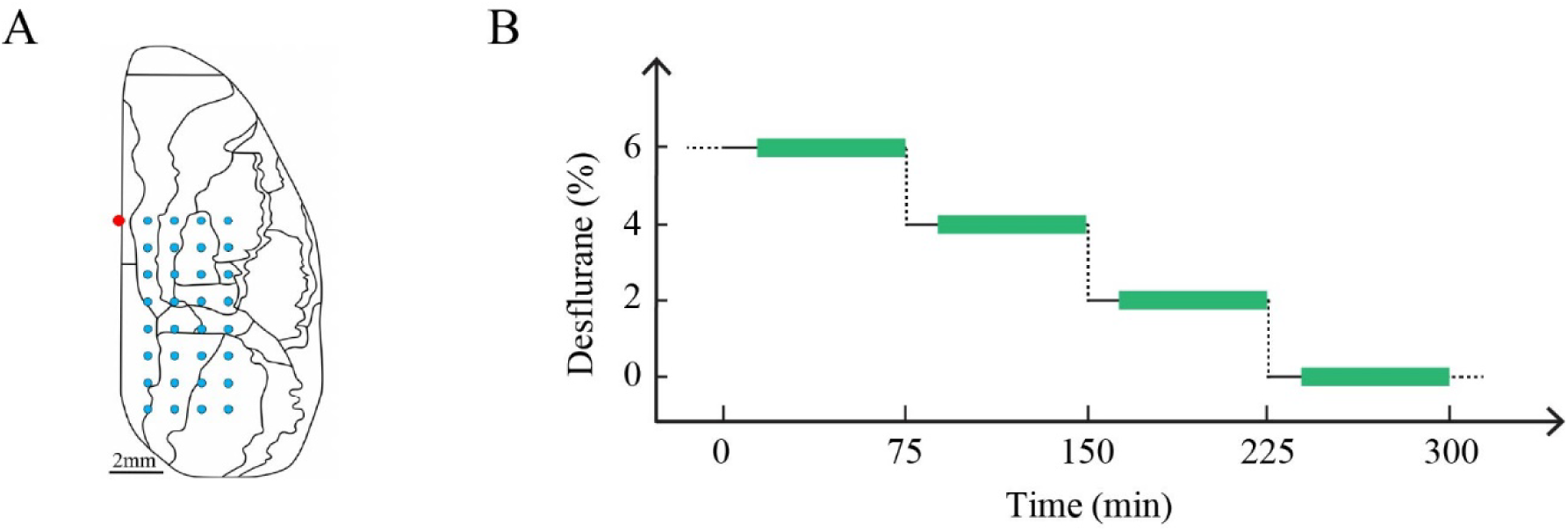
Schematic of electrocorticography recording and anesthetic protocol. **A**. A 32-site flexible polymer electrode array (4×8 grid) was chronically implanted in the right hemisphere of the rat. Red dot indicates location of Bregma. **B**. Desflurane was administered in reductions of steady-state concentrations. After a 15-min equilibration at each level, spontaneous electrocorticographic activity was recorded for 1 hour (green bars).

### Electrophysiological recording and preprocessing

Electrocorticographic signals were recorded at 1 kHz sampling rate and exported into MATLAB (version 2024b; MathWorks, Inc., Natick, MA). Four experiments were excluded due to excessive electrical noise, leaving 14 experiments for analysis (two experiments from each of six rats, and one experiment from each of the remaining two rats). Signals were first detrended using a local linear regression method with a 10-s window and 5-s step size (Chronux toolbox (Mitra and Bokil 2007)), and then lowpass filtered at 55 Hz using a 5^th^ order Butterworth filter applied with zero-phase forward and reverse algorithm. Channels with poor signal quality were identified visually and interpolated using the average of up to four nearest neighbors. Across experiments, 0-3 channels were interpolated. After interpolation, signals were re-referenced to the average. Data segments ± 1s around timestamps with amplitude exceeding the mean plus seven standard deviations in at least two channels were rejected. The percentage of data rejected was 0.41 ± 0.89 %, 0.40 ± 0.72 %, 4.09 ± 4.85 %, 7.56 ± 12.34 at 6 %, 4 %, 2 % and 0 % desflurane, respectively.

### Classification of dynamic states

To identify dynamic states, electrocorticographic signals from all four anesthetic concentrations were concatenated within each experiment. The classification procedure consisted of three sequential steps: feature extraction, within-experiment clustering, and cross-experiment alignment.

First, power spectrograms of signals from each channel were estimated using the multitaper method (Chronux toolbox (Mitra and Bokil 2007)) with a 2-s window, 50% overlap, a time-bandwidth product of 2, and 3 tapers, followed by smoothing with a moving average across 30 windows. This smoothing balanced temporal resolution, ensuring clusters were not dominated by brief fluctuations (seconds) or by long-timescale spectral properties of each anesthetic concentration (minutes).

Electrocorticographic spectrograms from all channels were concatenated and subjected to principal component analysis where principal components (PCs) capture the dominant variance of the data (channels × frequencies, 0-55 Hz with 0.5 Hz resolution). The spectrograms were projected into an N_PC_ -dimensional space, with N_PC_ chosen to maximize cluster separation across all anesthetic concentrations. Density-based clustering (Rodriguez and Laio 2014, Duan 2015) was then applied to the retained PCs. This method does not require predefining the number of clusters; cluster centers emerge naturally as points with locally high density and large distance from other high-density points.

After identifying clusters within each experiment, we aligned them to define a single set of canonical states that were uniform across all experiments. The final number and identity of the canonical states were determined through an interpretive review of the clusters obtained from individual experiments, identifying shared spectral and temporal features across them. The states were then ordered based on their predominant occurrence as the anesthetic concentration increased and their corresponding changes in delta power. To ensure cross-experiment consistency, individual clusters from each experiment were mapped onto the defined canonical states based on their proximity in the PC feature space. As demonstrated in our prior study (Li and Hudetz 2025), aggregating data at the cluster level, rather than the sample level, ensured an unbiased classification, preventing a single experiment’s long-duration state from disproportionately influencing the global state definition.

### Characterization of electrocorticographic properties across cortical states

State-specific electrocorticographic properties were characterized by using band-specific spectral power and spatiotemporal complexity. For each channel, normalized power spectra were obtained by dividing the absolute power by the total power from 0 to 55 Hz and then averaged across channels. Band-specific power values were extracted in delta (0-4 Hz), theta (4-8 Hz), and gamma (25-55 Hz) frequency bands. These ranges were selected for the primary analysis because they showed most robust power changes, clearly distinguishing states across experiments. Supplementary analyses examined state-dependent changes in higher gamma bands, extending up to 175 Hz.

Spatiotemporal complexity was quantified using the Lempel-Ziv complexity (LZC) algorithm (Lempel and Ziv 1976) and its normalized version to account for spectral content (Li and Mashour 2019). For each channel, the instantaneous amplitude was estimated using the Hilbert transform and binarized using the channel-specific mean as a threshold. Each epoch was then converted into a binary matrix, with rows representing channels and columns representing time points. LZC was calculated by sequentially scanning the matrix and counts the number of unique spatial patterns. To standardize the raw LZC values, they were scaled between 0 and 1 by dividing the raw LZC value by the mean of N = 50 surrogate data generated by randomly shuffling the temporal order of each epoch within each channel (representing a maximal complexity sequence). To isolate the complexity independent of spectral power, we then computed the normalized LZC (LZC_N_). This involves a second step of normalization where the scaled LZC value was divided by the mean LZC of N=50 surrogate series generated using the iterated amplitude-adjusted Fourier transform algorithm. The algorithm preserves the spectral profile of the original signal while randomizing phase. LZC_N_ values near 1 indicate complexity is fully explained by spectral content, whereas deviations from 1 indicate additional spatiotemporal dynamics.

### Quantification of cortical state occurrence across anesthetic concentrations

For each experiment, the state time series represents the temporal progression of cortical states across anesthetic concentrations. To characterize state occurrence and its concentration-dependency, we performed two complementary analyses: state occupancy at each concentration and concentration specificity of each state. For each anesthetic concentration, the fractional occupancy ratio was calculated as the proportion of time spent in each state. The degree of state co-occurrence at each concentration was quantified using Shannon entropy applied to the fractional occupancy ratios of all states, where an entropy of 0 indicates that only a single state was observed, and higher values indicate the coexistence of multiple states. For each cortical state, its occurrence rate across all concentrations was computed to evaluate concentration specificity. Shannon entropy quantified how restricted or widespread a state was across concentrations, with 0 indicating presence at only one concentration and higher values indicating presence across multiple concentrations.

To assess consistency across experiments, we quantified between-experiment dissimilarity in (i) state occupancy ratios (proportion of time in each state) at each concentration and (ii) concentration profiles (distribution of a state’s occurrence across concentrations) for each state. Dissimilarity for both was defined as the mean absolute difference between corresponding bins of two distributions, divided by 2 to constrain values between 0 and 1. This analysis evaluates whether cortical states exhibit similar concentration dependence across experiments.

### Analysis of temporal dynamics of cortical states

To characterize the temporal dynamics of cortical states, we assessed the stability of each state and the probabilities of state transitions across anesthetic concentrations. State stability was quantified using two measures. Dwell time was defined as the duration spent in a state before switching to a different state. Persistence likelihood was defined as the probability that the system remains in a given state at the next time step, equivalent to the fraction of time spent continuously in state *i*, *P*(*S*_*t*+1_ = *i* |*S*_*t*_ = *i*). For state transitions, we analyzed the sequence of switches by removing periods of state persistence from the state time series, counting only actual transitions between distinct states. The state transition matrix quantifies the probability of transitioning from state *i* to state *j*, with off-diagonal elements representing the probabilities of switching to a different state and all probabilities summing to 1 (Li, Vlisides et al. 2019). To examine the evolution of states during anesthetic emergence, we quantified transition asymmetry, defined as the difference between the probability of transition from state *i* to *j* and from *j* to *i*, reflecting directional preferences in the cortical state trajectory.

### Statistical analysis

Linear mixed model (LMM) analyses were performed using IBM SPSS Statistics version 30.0 for Windows (IBM Corp. Armonk, NY) for the following comparisons: (1) state-dependent electrocorticographic properties or state dynamic metrics across experiments, with state as a fixed effect and a random intercept for each experiment to account for inter-experiment variability; (2) concentration-dependent state dynamic metrics across experiments, with desflurane concentration as a fixed effect and a random intercept for each experiment; (3) between-experiment dissimilarity across states or concentrations, with state or concentration as a fixed effect and a random intercept for each pair of experiments; additionally, whether two experiments were performed in the same rat or in different rats was included as a fixed factor. All models were fitted using restricted maximum likelihood estimation. Post-hoc pairwise comparisons were performed using Student’s two-tailed paired t-tests, with Bonferroni correction applied for multiple comparisons.

To assess the spontaneous occurrence of cortical states across desflurane concentrations, Wilcoxon signed-rank tests were used to evaluate whether the entropy of state occupancy ratios at each concentration and the entropy of concentration profiles for each state differed from 0 (i.e., one-to-one correspondence between states and concentrations), with Bonferroni correction applied for multiple comparisons. Surrogate data analysis was performed to determine whether observed state transitions could occur by chance (Li et al., 2019). N = 1,000 surrogate time series were generated by randomly permuting the temporal order of state switches, while maintaining each state’s occurrence rate across experiments. For each surrogate time series, the transition probability was calculated for each pair of states, and a state transition was considered statistically significant if the original transition probability exceeded surrogate distribution. Across experiments, Wilcoxon signed rank test was used to compare state probabilities in the two directions for each pair of states to detect significant asymmetry. False discovery rate-adjusted *p* < 0.05 was used as the significance threshold. All the analyses were performed using MATLAB, unless otherwise specified. Normality of all datasets was assessed using Lilliefors-corrected Kolmogorov-Smirnov tests, and a *p*-value < 0.05 was considered statistically significant.

## Results

### Spectral properties distinguish cortical states

Spontaneous electrocorticographic signals and their power spectrograms exhibited distinct patterns during stepwise reduction of anesthetic concentrations (**Figure 2A**). To identify recuring cortical states, we performed principal component analysis on concatenated electrocorticographic spectrograms from all concentrations followed by density-based clustering of the first 3-7 principal components (PCs), which explained 66.2-86.0% of the variance (**Table S1**). This unsupervised procedure yielded 6 to 11 distinct clusters depending on the experiment (**Figure 2B**).

**Figure 2.**
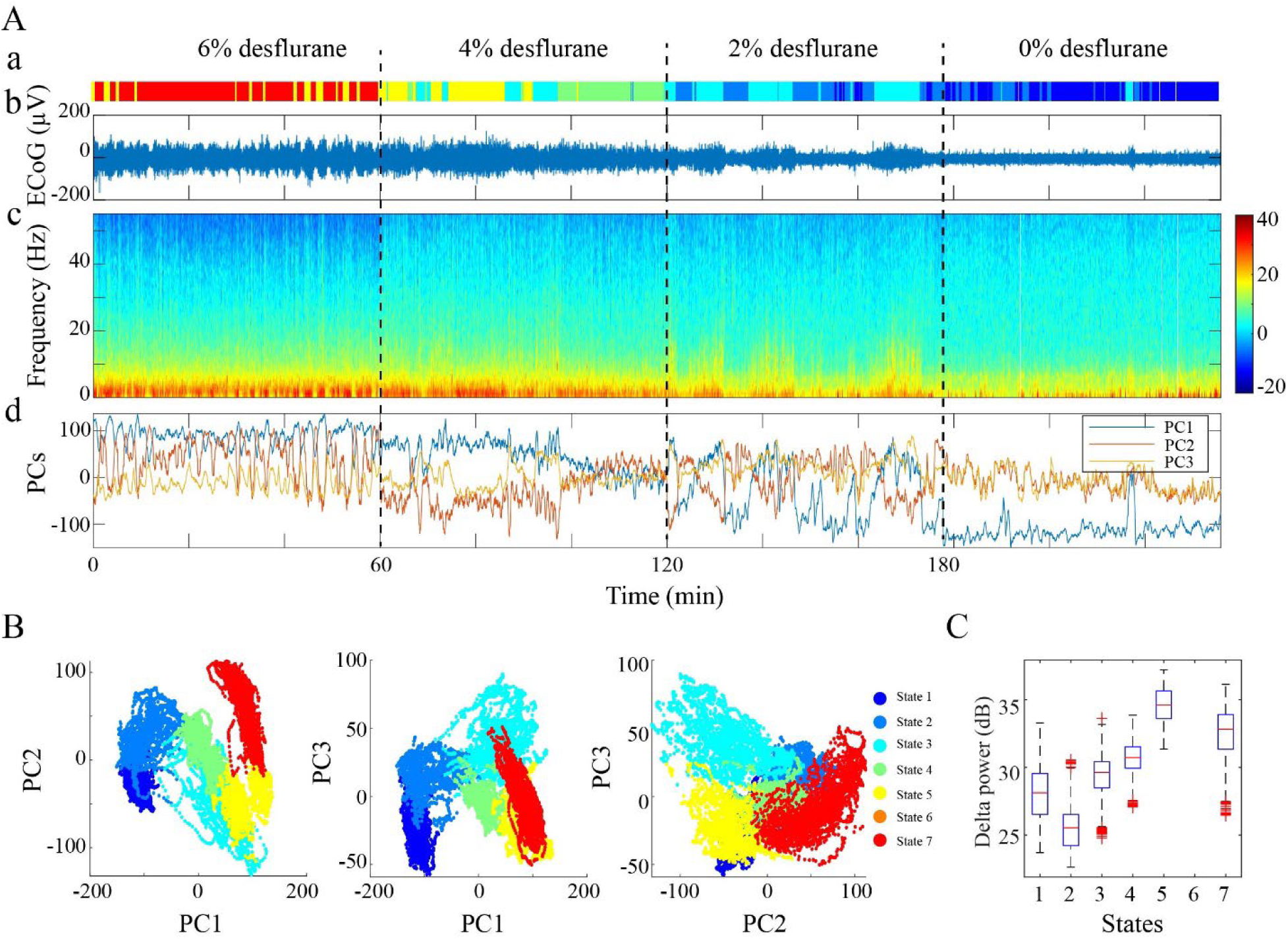
Classification of cortical states in a representative experiment. **A**. Seven states identified from principal component analysis and density-based clustering of electrocorticographic power spectrograms: (**a**) color-coded state labels, (**b**) example electrocorticographic signal from an occipital channel, (**c**) average power spectrogram across recording channels, (**d**) time course of first three principal components retained for density-based clustering. **B**. Pairwise scatterplots of the three principal components. Each point represents data from a 2-s time window with 1-s overlap. **C**. State-dependent delta power across all time windows within each state. Box plots indicate median and interquartile range, with whiskers extending to the most extreme values, and red crosses indicating outliers. ECoG, electrocorticography; PC, principal components.

For group-level analysis, clusters from individual experiments were mapped onto a unified set of seven canonical states (**Figure S1**). This consolidation was achieved through an interpretive review of the clusters from individual experiments, which revealed seven robust, recuring patterns that were consistent across the experimental cohort. The canonical states were then ordered according to their predominant occurrence across increasing anesthetic concentrations and their distinguishing spectral properties, particularly delta power (**Figure 2C**). In the PC feature space, the distribution of cortical states formed a consistent trajectory across experiments that aligned with the anesthesia depth and were arbitrarily designated as awake (State 1), light (States 2 and 3), intermediate (State 4), deep (State 5), and burst suppression (State 6). However, the precise trajectory varied between experiments, particularly in the positioning of the paradoxical State 7, which predominated at deep anesthesia but deviated from the general trajectory shaped by States 1-6 (**Figure S2**).

The seven cortical states exhibited distinct electrocorticographic properties (**Figure 3A-B**). State 1, which occurred predominant at 0% desflurane showed high theta and gamma activity (**Figure 3 C**). From State 1 to State 5, theta and gamma activity decreased, while delta power increased (**Figure 3B, D**). States 5, 6 and 7 were predominantly observed at 6% desflurane. State 6 marked the onset of burst suppression, with a suppression ratio of 15.39[11.35, 23.73] % (median [IQR]) (**Figure 3E**). State 7 (paradoxical state) diverged from this trend, showing significantly lower delta power (*p*<0.001) and elevated theta (*p*=0.005) and gamma power (*p*=0.002) relative to State 5 (**Figure 3D, Table S2**). High-gamma power showed similar state-dependent changes (**Figure S3**), whereas electromyographic power did not appear to increase in State 7 (**Figure S4**).

**Figure 3.**
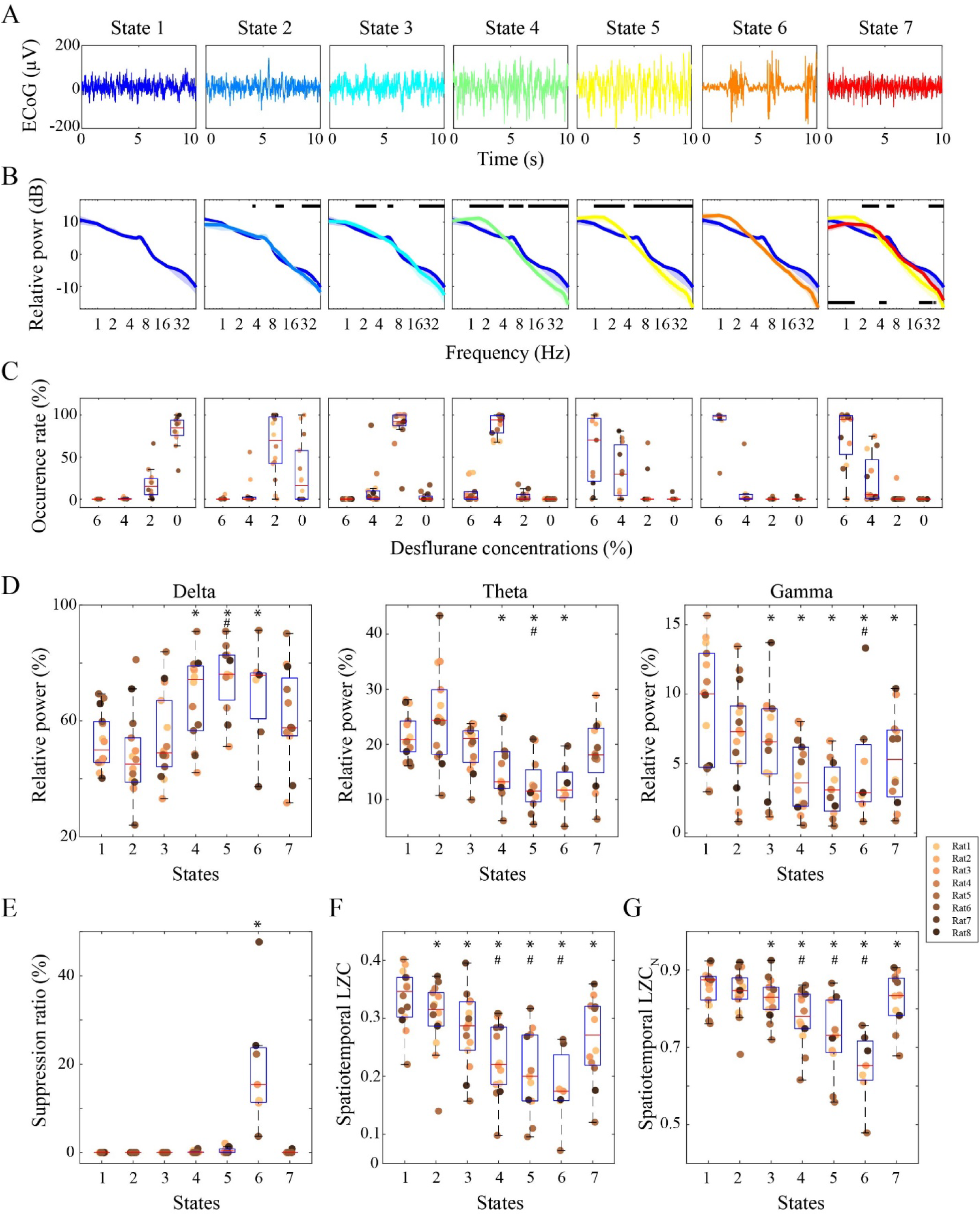
Electrocorticographic properties of cortical states. **A**. Example electrocorticographic traces from an occipital channel of one experiment. **B**. Power spectra normalized to total power (0-55 Hz) and averaged across all channels from all experiments. Colored lines and shaded areas represent the median and interquartile range of power density across experiments. For comparison, State 1 spectrum was replotted in all other states, and the State 5 spectrum was included in the plot for State 7. In States 2-7, horizontal black bars at the top indicate frequency ranges that differ significantly from State 1; in State 7, horizontal black bars at the bottom indicate difference from State 5 (false discovery rate-adjusted *p*<0.05, Wilcoxon signed-rank test). **C**. State occurrence rates across desflurane concentrations. **D**. State-dependent changes in delta, theta and gamma power. **E**. Changes in suppression ratio across states. **F, G**. State-dependent changes in spatiotemporal complexity as quantified by the Lempel-Ziv complexity (LZC) algorithm (**F**) and by normalized LZC relative to surrogate data generated by the iterative amplitude-adjusted Fourier transform algorithm (**G**). In panels **C**-**G**, box plots show the median and interquartile range, whiskers extend to the most extreme values, and data points from individual experiments are shown as dots. Two recording sessions were performed for Rats 1-6. * *p*<0.05/6 vs. State 1. # *p*<0.05/6 vs. State 7, linear mixed models. ECoG, electrocorticography; LZC, Lempel-Ziv complexity.

### State-dependent changes in electrocorticographic signal complexity

Beyond channel-averaged spectral features, we investigated the spatiotemporal complexity of electrocorticographic signals across cortical states using Lempel-Ziv complexity (LZC), which quantifies the number of unique patterns across channels and time points and captures variability of cortical activity both spatially and temporally. LZC decreased progressively from State 1 to State 6 (all *p*≤0.005 vs. State 1; **Figure 3F**). State 7 partially reversed this trend, showing a significant increase relative to States 4-6 (all *p*≤0.005), although its complexity remained lower than that of State 1 (*p*<0.001).

To determine whether these changes were solely driven by state-dependent spectral differences, we computed normalized LZC (LZC_N_) by dividing each observed LZC value by the mean LZC obtained from surrogate data that preserved the original spectral power but randomized phase relationships. LZC_N_ minimizes the predictable components captured by the power spectrum and isolates complexity arising from the unpredictable components of the temporal structure and statistical dependencies of the signal. LZC_N_ closely mirrored the LZC trends. Compared with State 1, LZC_N_ decreased in all states (all *p*≤0.005) except State 2 (*p*=0.154). Importantly, LZC_N_ changes were non-monotonic: State 7 exhibited significantly higher complexity values than States 4-6 (all *p*<0.001) (**Figure 3G** and **Table S2**), suggesting a rebound in complexity beyond what is reflected by spectral power.

### Spontaneous occurrence of cortical states across desflurane concentrations

Having characterized the electrocorticographic properties of each state, we next examined their occurrence and concentration-dependency across desflurane levels. **Figure 4A** shows the temporal sequences of cortical states in individual experiments during stepwise reductions of anesthesia. While certain states predominated at specific concentration (e.g. State 4 at 4%, States 5-7 at 6%), other states also appeared at the same concentration. For example, State 4 was observed at 6% desflurane in 8 of 14 experiments (**Figure 4 A, B**).

**Figure 4.**
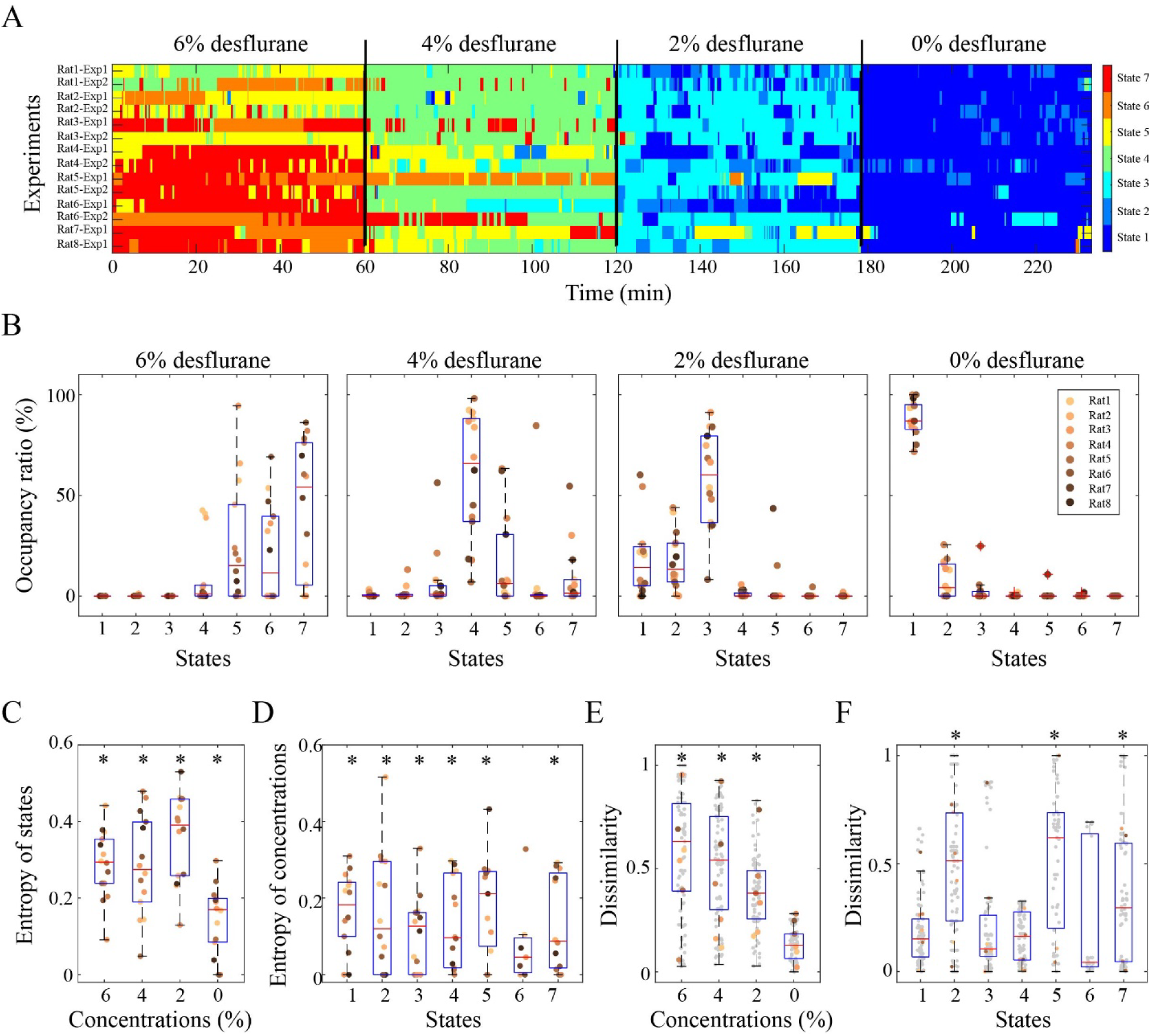
Spontaneous occurrence of cortical states across desflurane concentrations. **A**. Temporal progression of cortical states in each experiment during stepwise reduction of desflurane concentration. For visualization, the duration at each desflurane concentration was rescaled to the median across all experiments. **B**. Occupancy ratio, defined as the proportion of time spent in each state at a given anesthetic concentration, across all experiments. **C**. Entropy of state occupancy ratios, quantifying the degree of state co-occurrence at each concentration across all experiments. * significantly greater than 0 (*p*<0.05/4, Wilcoxon signed-rank test), reflecting the presence of multiple coexisting states. **D**. Entropy of the occurrence rates of a given state, quantifying the degree of its occurrence across multiple concentrations. * significantly greater than 0 (*p*<0.05/7, Wilcoxon signed-rank test), reflecting that the state is present across multiple concentrations. **E**. Between-experiment dissimilarity of state occupancy ratios at each concentration. * *p*<0.05/3 vs. 0% desflurane, linear mixed models. **F**. Between-experiment dissimilarity of the concentration profile for each state. * *p*<0.05/6 vs. State 1, linear mixed models. In panels **B**-**D**, dots represent single experiments; in panels **E**-**F**, dots indicate pairs of experiments, with colored dots for pairs from the same rat and gray dots for pairs from different rats.

To quantify state co-occurrence, we calculated the Shannon entropy of state occupancy ratios for each concentration, which measures the diversity of states present. Entropy values were 0.17 [0.08, 0.20] at 0%, 0.40 [0.26, 0.46] at 2%, 0.27 [0.19, 0.40] at 4%, and 0.29 [0.24, 0.35] at 6% desflurane — all significantly greater than 0, indicating that no concentration was occupied by a single state (**Figure 4C**). Conversely, each state occurred across multiple concentrations (**Figure 3C**), and concentration specificity was quantified using the entropy of each state’s occurrence across all concentrations. Entropy values were significantly greater than 0 for all states (*p*<0.001), except for the burst suppression (State 6; **Figure 4D**). Together, these results indicate that cortical states emerged spontaneously and were not strictly determined by desflurane concentration.

To evaluate whether cortical states showed consistent concentration dependency across experiments, we compared the proportion of time each state occupied at each anesthetic level (state occupancy ratio) and how each state’s occurrence was distributed across concentrations (concentration profile). Comparisons between experiments from the same rat (within-animal variability) to those from different rats (between-animal variability) revealed no significant difference for either measure (occupancy ratio: F(1,170.526) = 0.281, *p* = 0.597; concentration profiles: F(1,237.539) = 0.298, *p* = 0.586; **Figure 4E, F**), indicating that inter-animal differences did not contribute substantially to the variability. Instead, variability across experiments was primarily determined by concentration and state identity. Between-experiment dissimilarity of state occupancy ratios increased with anesthetic concentration: 0.13 [0.07, 0.18] at 0%, 0.38 [0.26, 0.49] at 2%, 0.54 [0.30, 0.75] at 4% and 0.63 [0.39, 0.82] at 6% desflurane (all *p*<0.001; **Figure 4E**), suggesting that the patterns of state occupancy became less consistent across animals as anesthesia deepened. Dissimilarity of concentration distributions varied by state: it was low for State 1, significantly higher for States 2, 5, and 7 (all *p*<0.001), and comparable to State 1 for the remaining states (**Figure 4F**).

### Temporal dynamics of cortical states across desflurane concentrations

Following the characterization of state occurrence, we next examined the temporal organization of cortical states to reveal how they persist and transition across anesthetic concentrations and how they evolve during anesthetic emergence.

We first quantified how long each state persisted and how likely it was to switch to another state. Across all states, mean dwell time was 136.55 (76.70, 217.85) s. Although dwell times differed moderately across states (F(6,79) = 2.552, *p*=0.026), post-hoc comparisons were not significant after correction for multiple testing (**Figure 5A**). Overall, states were highly persistent, with a likelihood of remaining in the same state of 99.36 (98.81, 99.62) %. States 2 and 5 showed slightly lower persistence, whereas the other states were comparable to State 1 (**Figure 5B**).

**Figure 5.**
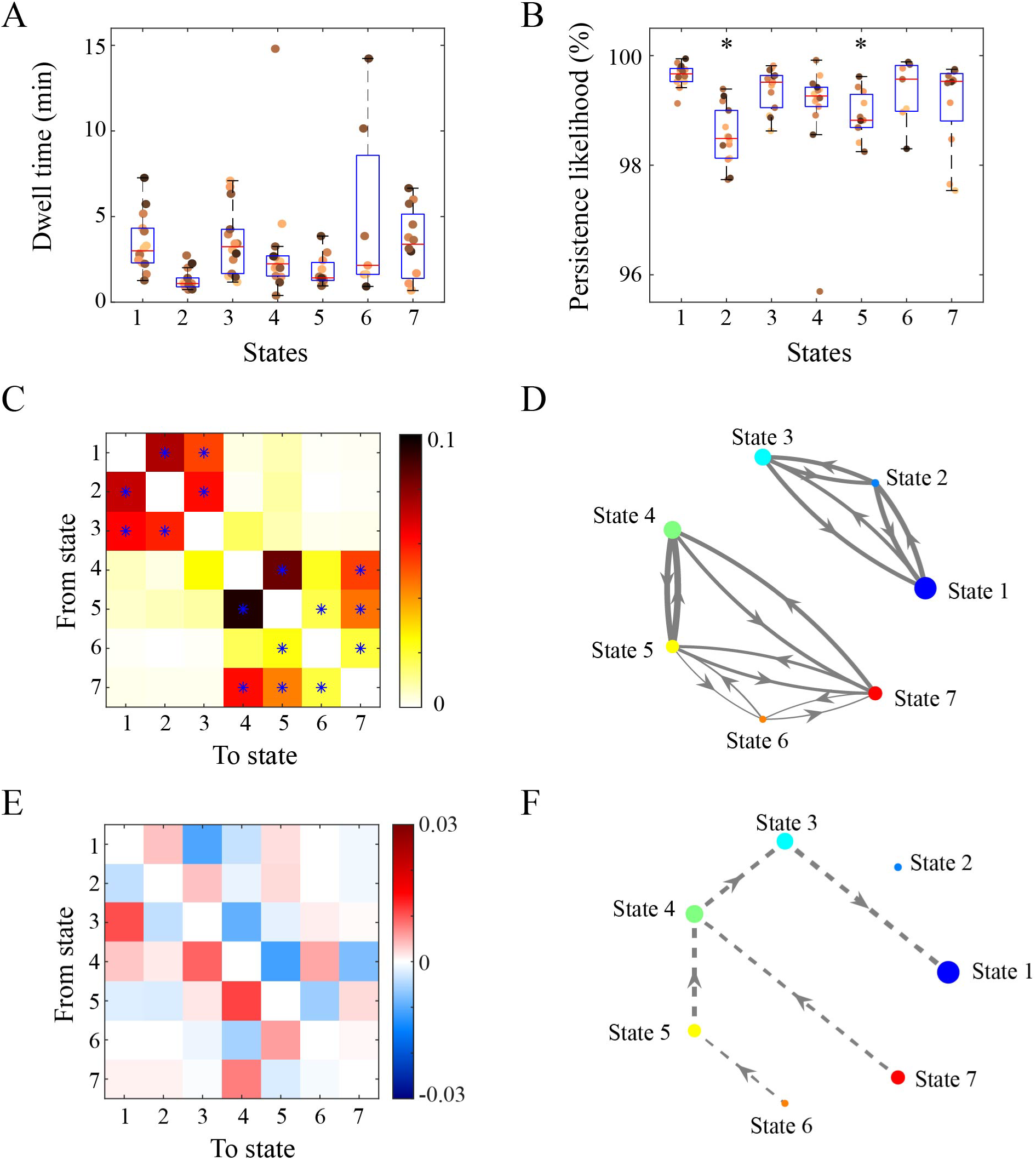
State transition dynamics across desflurane concentrations. **A**. Averaged dwell time of each state across all experiments. **B**. Likelihood of state persistence across experiments. * *p*<0.05/6 vs. State 1, linear mixed models. **C**. State transition probability matrix from pooled data across all experiments. Off-diagonal elements represent the probability of transitioning from a state (row) to another state (column). * probabilities significantly higher than expected by chance, based on surrogate data generated by random permutation of state time series (false discovery rate-adjusted *p* < 0.05). **D** Graphical representation of significant state transitions. Graph nodes represent states, node size is proportional to persistence probability, and directed, weighted edges indicate transition probabilities between states. **E**. Asymmetry of state transitions. Each element represents the difference between the transition probability from state *i* to *j* and the transition probability in the opposite direction, from *j* to *i*, averaged across experiments. Red elements indicate transitions that are more likely than the opposite direction (blue elements). **F**. State evolution during stepwise anesthetic reduction. The top 25% of transitions with the highest probability difference as identified from E are shown.

We next analyzed between-state transition probabilities to determine whether transitions occurred randomly or followed structured pathways. The group-level transition probability matrix, compared against surrogate data preserving state occurrence rates, revealed that certain transitions occurred more frequently than expected by chance (FDR-corrected *p* < 0.05; **Figure 5C, D**). In the transition matrix two major clusters emerged. The first cluster involved states predominantly associated with awake to light anesthesia (States 1-3), which transitioned frequently among one another. The second cluster comprised states primarily associated with intermediate to deep anesthesia (States 4-7). Within this cluster, transitions among states predominant at 6% desflurane were more frequent than expected. Switches between the intermediate anesthesia state (State 4) and deep/paradoxical states (States 5 and 7) were also significant, whereas transitions between State 4 and the burst suppression state (State 6) did not exceed chance probability. These results indicate that cortical state transitions are structured and hierarchical: the brain preferentially moves between intermediate and less extreme deep states (States 5 and 7) before reaching the most suppressed state (State 6), with frequent transitions among deep states 5-7 indicating organized exploration within the deep anesthesia cluster. Moreover, transitions between the two clusters were not significant, suggesting that the cortical state space operates in two distinct regimes—light and deep anesthesia—separated by a strong resistance to transition between them.

We also tested whether transitions were asymmetric to identify preferred trajectories embedded in cortical state dynamics during anesthetic emergence. Across experiments, transitions exhibited a modest directional preference from deeper to lighter anesthesia (**Figure 5 D, E**). Burst suppression (State 6) tended to transition to deep anesthesia (State 5), and together with the paradoxical state 7, preferentially transitioned to the intermediate anesthesia state (State 4). State 4, in turn, tended to transition to lighter state (State 3) and ultimately the awake state (State 1). The strongest directional bias was from State 4 to State 3 (*p* = 0.0039), although this did not remain significant after correcting for multiple comparisons. This overall directional flow towards lighter states is consistent with the experimental protocol of stepwise anesthetic reduction. However, it also reveals the brain’s intrinsic dynamic organization: cortical state dynamics follow a specific trajectory when transitioning from deep to light anesthesia regime during emergence from desflurane anesthesia.

Collectively, these results demonstrate that it is more probable for the brain to remain in a given state than to switch to another, but when transitions occur, they follow structured and non-random pathways shaped by both intrinsic state dynamics and externally driven anesthetic reduction.

## Discussion

To determine how large-scale electrocortical state transitions relate to anesthetic concentration, we analyzed hemispheric electrocorticographic signals in rats during a protocol spanning three sustained concentrations of desflurane and wakefulness. Using unsupervised clustering of electrocorticographic power spectrograms, we identified a consistent set of seven discrete states characterized by distinct spatiotemporal complexity. Six states followed the anesthetic concentration although the states’ occurrence was not strictly tied to any anesthetic level. A notable exception was a paradoxical state, which emerged predominantly during deep anesthesia while exhibiting unexpectedly low delta power and increased complexity—features indicative of an organized neural state. Transition analysis of controlled emergence from anesthesia further revealed structured, non-random pathways between states, primarily confined to light- or deep-anesthesia regimes, with a mild directional bias from deep to light states consistent with recovery of consciousness. Together, these results indicate that electrocortical activity under desflurane involves spontaneous dynamics and paradoxically activated states with increased global complexity during deep anesthesia.

Consistent with a growing body of prior evidence (Hudson, Calderon et al. 2014, Li, Vlisides et al. 2019, Lee, Wang et al. 2020, Li and Hudetz 2025), our findings demonstrated that different cortical states emerge spontaneously and exhibit a many-to-many relationship with various clinically relevant anesthetic concentrations. The variability of cortical states at each anesthetic level was reliably present in all experiments, although the number and distribution of states varied among the experiments. Both the within-experiment diversity of states and the between-experiment inconsistency of state-concentration relationships were amplified at deeper anesthetic levels suggesting that the anesthetic enhanced stochastic state fluctuations and weakened the deterministic coupling between cortical states and anesthetic concentration. Moreover, within-animal variability was comparable to between-animal variability, mirroring previous observations in genetically identical mice that stochastic state switching—rather than inter-individual differences—underlies variability in emergence from general anesthesia (Stone, Kelz et al. 2025). While the spontaneous nature of state fluctuations and their temporal dynamics align with recent reports of multiple trajectories of cortical activity under anesthesia (Curic, Ashby et al. 2024), our results provide a critical reinterpretation of earlier inconsistences. Specifically, awake-like states observed at surgical levels of isoflurane or ketamine anesthesia—previously attributed to inter-subject variability within a unitary framework (Curic, Ashby et al. 2024)—are more plausibly intrinsic manifestations of spontaneous state transitions occurring within individual animals.

Our state transition analysis also revealed a metastable organization of cortical dynamics (Hancock, Rosas et al. 2025) during anesthesia. The cortex preferentially transitioned between two clusters of states—one corresponding to presumably conscious lightly sedated or awake condition and the other to arguably unconscious intermediate or deep anesthesia—indicating that the anesthetized brain favors to operate within two discrete dynamic regimes. Transitions between these regions were rare, suggesting strong attractor boundaries that stabilize each regime. Only when the anesthetic concentration was sufficiently low, did the brain exit the “deep” cluster and re-enter the “light” cluster, where it again explored multiple states within that regime. These structured yet concentration-related transitions suggest that anesthetic depth governs not only the overall level of cortical suppression but also the accessibility and stability of distinct neural states, reflecting an interaction between intrinsic neural organization and external pharmacologic drive. Moreover, the structured and discontinuous progression of cortical states during emergence aligns with previous observations of non-continuous recovery pathways under general anesthesia (Chander, García et al. 2014, Hudson, Calderon et al. 2014).

The discovery of paradoxical state in electrocorticographic activity is important. This essentially global state was characterized by low delta power and an unexpected rebound of complexity during deep anesthesia. This finding is at variance with our former report of paradoxical state in visual cortex unit activity, which indicated high and asynchronous firing but low complexity (Lee, Wang et al. 2020, Li and Hudetz 2025). However, the two states in question may not be equivalent. Global paradoxical state from electrocorticography analysis was associated with low electromyographic power, whereas the local paradoxical state from spike data was accompanied by high electromyographic power (Lee, Wang et al. 2020) suggesting that they reflected distinct neurophysiological conditions. Indeed, local states can be dissociated from global states: for example, local activation in motor cortex occurs during sleep (Nobili, Ferrara et al. 2011), and wake-like activity in layer 5 cortex can be induced and maintained under deep anesthesia (Pardo-Valencia, Moreno-Gomez et al. 2024), highlighting a complex interplay between global and local brain states (Cooke and Balbinot 2024). Moreover, recent evidence suggests that global state transitions are shaped by a multitude of local cortical activity patterns that are only weakly correlated across local regions (Blackwood, Shortal et al. 2022). Within this framework, the elevated global complexity of the paradoxical state may not reflect uniformly high complexity but rather increased functional heterogeneity and diversity across cortical regions.

The elevated global complexity may indicate that, despite the strong suppression of baseline activity, a certain degree of residual information integration persists. Unlike rapid eye movement sleep, where high electromyographic accompanies global high complexity (Cavelli, Mao et al. 2023), the present state shows low electromyographic power, indicating minimal behavioral activation. The observed paradoxical state during deep anesthesia may reflect the working of a compensatory mechanism—a dynamical attempt by the cortex to counterbalance the profoundly depressed states induced by high desflurane concentrations (State 5 and burst suppression). Although high complexity alone does not constitute evidence of consciousness, the emergence of this state bears resemblance to the alternating low- and high-complexity patterns previously observed under ketamine anesthesia (Li and Mashour 2019), where slow delta oscillations (low complexity) alternate with gamma oscillations (high complexity) in the so-called gamma-burst pattern. While this alternation may be related to the distinctive pharmacological profile of ketamine, which has both anesthesia and psychedelic properties, we cannot exclude the theoretical possibility that the present state under desflurane reflects a reorganized cortical network capable of limited signal processing. The persistence of high complexity may thus reflect residual, partial information integration, suggesting that the anesthetized brain retains some capacity for structured neural dynamics, and perhaps fragments of consciousness, even under profound activity suppression.

The present findings also have important implications for anesthetic practice. The many-to-many relationship between anesthetic concentration and cortical state, including the emergence of high-complexity paradoxical state during deep anesthesia, poses a major challenge for conventional dose-based monitoring. For example, as paradoxical State 7 exhibits spectral and complexity features resembling lighter planes of anesthesia, standard monitors may misinterpret it as a sign of arousal, prompting clinicians to unnecessarily increase anesthetic delivery in pursuit of a deeper state and thereby risking overdose and related complications. Conversely, the appearance of a deep cortical state at a low concentration could lead to premature dose reduction and inadequate anesthesia or intraoperative awareness. These findings underscore the need for monitoring approaches that incorporate spontaneous state fluctuations and global cortical dynamics to improve dosing accuracy and patient safety. Moreover, the increased between-experiment variability observed at higher anesthetic depths highlights the importance of individualized monitoring. Finally, the structured nature of cortical state transitions and their metastable organization may offer new insights into the neural mechanisms of emergence, potentially enabling targeted modulation to accelerate or stabilize recovery.

The present study has several methodological limitations. First, to examine the relationship with anesthetic concentration, cortical states were classified on a relatively slow timescale using 2-s windows with 1-s overlap and smoothing over 30 windows; shorter windows may reveal additional cortical states at a finer grain. Second, the paradoxical state was operationally defined as any cortical pattern deviating from the expected monotonic relationship with anesthetic concentration. While exhibiting low delta power and high complexity during deep anesthesia, it may include heterogeneous subtypes—such as prolonged or transient states—that future work could further differentiate. Third, although spontaneous complexity aligned with perturbational complexity (Breyton, Fousek et al. 2025), approaches integrating perturbational paradigms may uncover properties of paradoxical state not captured by spectral or spontaneous complexity measures. Fourth, the absence of behavioral monitoring prevents excluding natural sleep that may occur at 2% and 0% desflurane. Fifth, hemispheric electrocorticogram recordings captures global but not local network dynamics, and multi-scale studies incorporating local field potential or unit recordings in the same subject would be needed to clarify local-global dissociations (Maria V. Sanchez-Vives 2025). Sixth, this study focused solely on desflurane; future work is necessary to determine whether spontaneous state transitions and paradoxical high-complexity state generalize across anesthetic agents of different pharmacological classes. Finally, while rat electrocorticographic recording reveals large-scale cortical organization, translation to humans should be cautious given the greater complexity of human cortical architecture, dynamics, and anesthetic responses.

In summary, cortical dynamics under desflurane anesthesia exhibit spontaneous transitions among multiple discrete states, including a paradoxical high-complexity state during deep anesthesia, and a structured, metastable organization rather than a monotonous concentration-dependent progression. These findings provide new insights into the neural mechanisms of anesthesia and may inform improved monitoring, dosing, and recovery strategies in clinical practice.

## Acknowledgements

Research reported in this publication was supported by the National Institute of General Medical Sciences of the National Institutes of Health under award number R01-GM056398 and the Center for Consciousness Science, Department of Anesthesiology, University of Michigan Medical School, Ann Arbor, Michigan, USA. The content is solely the responsibility of the authors and does not necessarily represent the official views of the National Institutes of Health. The authors express their gratitude to Dr. Shiyong Wang for his assistance in performing the experiments, to Drs. Gábor Juhász and Zsolt Borhegyi, Eötvös Loránd University, Budapest, Hungary and to Dr. Zoltán Fekete, Pázmány Péter Catholic University, Budapest for their advice for the use of flexible polymer electrodes.

